# Resection and repair of a Cas9 double-strand break at CTG trinucleotide repeats induces local and extensive chromosomal rearrangements

**DOI:** 10.1101/782268

**Authors:** Valentine Mosbach, David Viterbo, Stéphane Descorps-Declère, Lucie Poggi, Wilhelm Vaysse-Zinkhöfer, Guy-Franck Richard

**Affiliations:** Institut Pasteur, CNRS, UMR3525, 25 rue du Dr Roux, F-75015 Paris, France; Sorbonne Universités, Collège doctoral, F-75005 Paris, France; Biologics Research, Sanofi R&D, 13 Quai Jules Guesde, 94403 Vitry sur Seine, France; Institut Pasteur, Center of Bioinformatics, Biostatistics and Integrative Biology (C3BI), F-75015 Paris, France; Institut de Génétique et de Biologie Moléculaire et Cellulaire (IGBMC), UMR7104 CNRS/Unistra, INSERM U1258, 1, rue Laurent Fries BP 10142, 67404 Illkirch, France

**Keywords:** Trinucleotide repeats, CRISPR-Cas9, double-strand break resection, chromosomal rearrangements, gene conversion

## Abstract

Microsatellites are short tandem repeats, ubiquitous in all eukaryotes and represent ∼2% of the human genome. Among them, trinucleotide repeats are responsible for more than two dozen neurological and developmental disorders. Targeting microsatellites with dedicated DNA endonucleases could become a viable option for patients affected with dramatic neurodegenerative disorders. Here, we used the *Streptococcus pyogenes* Cas9 to induce a double-strand break within the expanded CTG repeat involved in myotonic dystrophy type 1, integrated in a yeast chromosome. Repair of this double-strand break generated unexpected large chromosomal rearrangements around the repeat tract. These rearrangements depended on *RAD52*, *DNL4* and *SAE2*, and both non-homologous end-joining and single-strand annealing pathways were involved. Resection and repair of the double-strand break (DSB) were totally abolished in a *rad50*Δ strain, whereas they were impaired in a *sae2*Δ mutant, only on the DSB end containing most of the repeat tract. This proved that Sae2 plays significant different roles in resecting a DSB end containing a repeated and structured sequence as compared to a non-repeated DSB end.

In addition, we also discovered that gene conversion was less efficient when the DSB could be repaired using a homologous template, suggesting that the trinucleotide repeat may interfer with gene conversion too. Altogether, these data show that *Sp*Cas9 is probably not a good choice when inducing a double-strand break at or near a microsatellite, especially in mammalian genomes that contain many more dispersed repeated elements than the yeast genome.

## Introduction

Microsatellites are short tandem repeats ubiquitously found in all eukaryotic genomes sequenced so far (Richard et al., 2008). Altogether, they cover ∼2% of the human genome, a figure similar to the whole protein-coding sequence (International Human Genome Sequencing Consortium, 2004). Naturally prone to frequent repeat length polymorphism, some microsatellites are also prone to large expansions that lead to human neurological or developmental disorders, such as trinucleotide repeats involved in Huntington disease, myotonic dystrophy type 1 (Steinert disease), fragile X syndrome or Friedreich ataxia (Orr and Zoghbi, 2007). These expansion-prone microsatellites share the common property to form secondary DNA structures *in vitro* (Gacy et al., 1995) and genetic evidences suggest that similar structures may also form *in vivo*, transiently stalling replication fork progression (Anand et al., 2012; Nguyen et al.; Pelletier et al., 2003; Samadashwily et al., 1997; Viterbo et al., 2016). Among those, CCG/CGG trinucleotide repeats are fragile sites in human cells, forming frequent double-strand breaks when the replication machinery is slowed down or impaired (Sutherland et al., 1998). Similarly, CAG/CTG and CCG/CGG microsatellites are also fragile sites in *Saccharomyces cerevisiae* cells (Balakumaran et al., 2000; Freudenreich et al., 1998). Therefore, microsatellite abundance and the natural fragility of some of them make these repeated sequences perfect targets to generate chromosomal rearrangements potentially leading to cancer.

Double-strand break (DSB) repair mechanisms have been studied for decades in model organisms as well as in human cells and led to the identification of the main genes involved in this process (reviewed in Krogh and Symington, 2004). Many of these advances were made possible by the use of highly specific DNA endonucleases, such as the meganucleases I-*Sce* I or HO (Fairhead and Dujon, 1993; Haber, 1995; Plessis et al., 1992). Other frequently used methods involved ionizing radiations making genome-wide DSBs (Nelms et al., 1998). However, the fate of a single double-strand break within a repeated and structured DNA sequence has never been addressed, until recently. In a former work, we used a TALE Nuclease (TALEN) to induce a unique DSB into a long CTG trinucleotide repeat integrated into a *S. cerevisiae* chromosome. We showed that 100% of yeast cells in which the TALEN was expressed exhibited a large contraction of the repeat tract, going from an initial length of ∼80 CTG triplets to less than 35. *POL32*, *DNL4* and *RAD51* were shown to play no detectable role in repairing this DSB. On the contrary, *RAD50*, *RAD52* and *SAE2* were required for proper repair of the DSB, and a functional Sae2 protein was found to be essential for efficient DSB resection, suggesting that repeat contraction occurred by a single-strand annealing (SSA) process, involving preliminary resection of the break, followed by annealing of the two DSB ends carrying the repeat tract (Mosbach et al., 2018; Richard et al., 2014).

In the present work, we used the *Streptococcus pyogenes* Cas9 endonuclease (*Sp*Cas9) to induce a DSB within the same long CTG trinucleotide repeat integrated in the yeast genome. The break was made at the 3’ end of the repeat tract (Figure 1A), using a guide RNA that targets the repeat tract. Frequent rearrangements were found in surviving cells, with local deletions as well as more extensive ones involving recombination between retrotransposon LTRs. Survival and repair depended on *RAD50*, *RAD52*, *SAE2* and *DNL4* and double-strand break resection was abolished in *rad50*Δ and *sae2*Δ mutants. A more specific version of the nuclease, Enhanced *Sp*Cas9, generated the same rerrangements. In addition, we also discovered that gene conversion was less efficient when *Sp*Cas9 was used to induce a DSB within a CTG repeat tract that could be repaired with a homologous template, suggesting that the trinucleotide repeat may interfer with gene conversion too.

**Figure 1:**
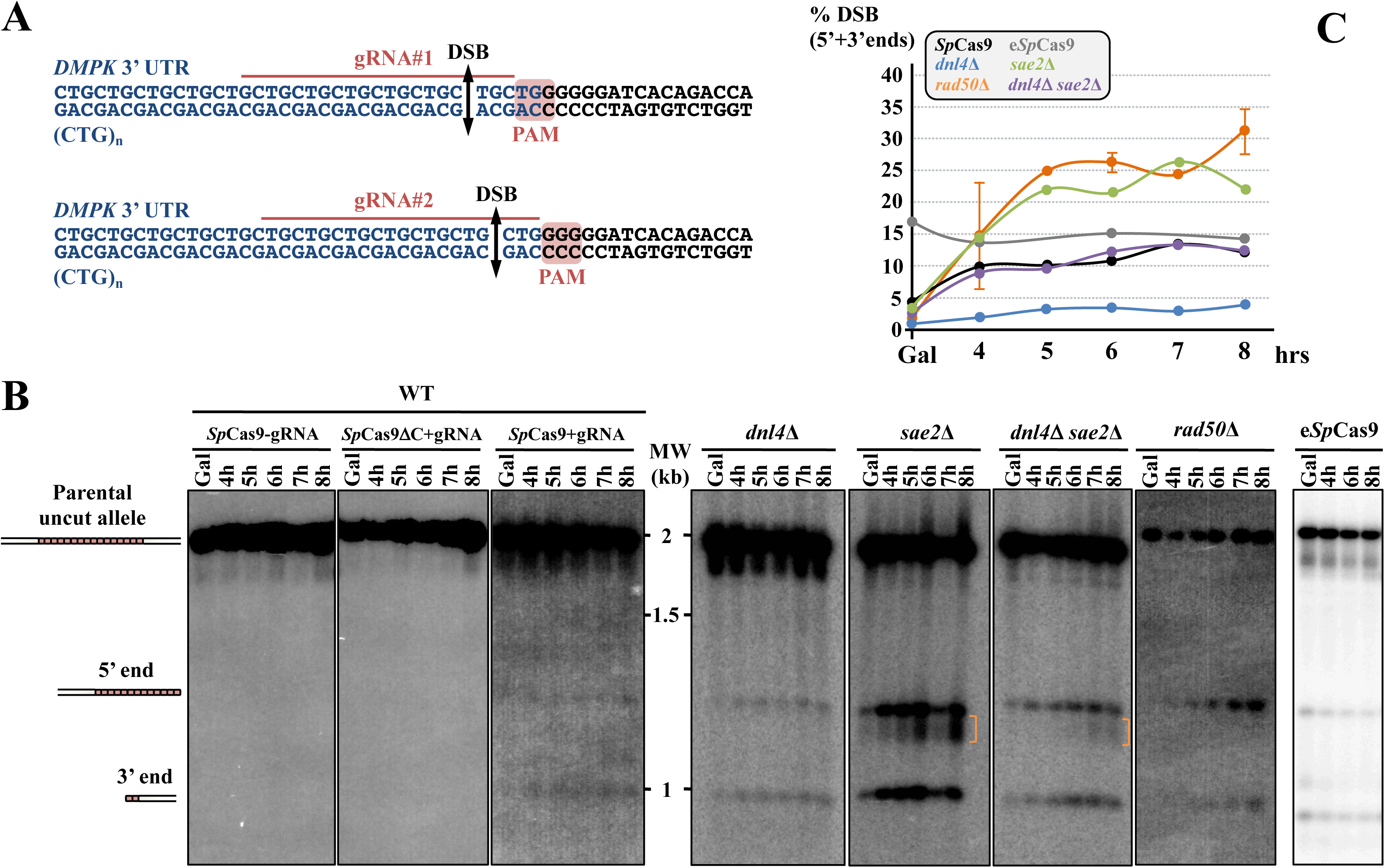
*Sp*Cas9 and e*Sp*Cas9 DSB induction in wild type and mutant strains. **A:** Sequence of the *SUP4*::(CTG)n locus. The CTG trinucleotide repeat tract comes from a human DM1 patient and is shown in blue. The flanking non-repeated DNA is in black. For each guide RNA, the PAM, the gRNA sequence as well as the expected DSB site are indicated. **B**: Southern blots of yeast strains during the time course. Lanes labeled *Sp*Cas9-gRNA and *Sp*Cas9ΔCter+gRNA are control strains in which no DSB was visible. In the strain expressing both *Sp*Cas9 and the gRNA, two bands are visible in addition to the parental allele (1966 bp). One band corresponds to the 3’ end of the DSB containing a small number of triplets (821 bp), the other one corresponds to the 5’ end of the DSB containing most of the repeat tract (1145 bp). **C:** Quantification of 5’ and 3’ DSB signals. For each time points, the total 5’ + 3’ signals were quantified and plotted as a ratio of the total signal in the lane. Three independent time courses were run in each strain background (except *rad50*Δ for which two time courses were run) and plots show the average of three (or two) time courses.

## Results

### A Cas9-induced double-strand break within CTG repeats induces cell death and chromosomal rearrangements around the repeat tract

In previous works, we showed that a TALEN targeted at the CTG trinucleotide repeat from the human *DMPK* gene 3’ UTR integrated in a yeast chromosome, was extremely efficient at contracting the repeat tract below the pathological length (Mosbach et al., 2018; Richard et al., 2014). In order to determine whether the CRISPR-Cas9 system could be used in the same manner, a plasmid-borne *Streptococcus pyogenes* Cas9 nuclease (*Sp*Cas9) was expressed in *Saccharomyces cerevisiae* from a *GAL1*-inducible promoter (DiCarlo et al., 2013). The same plasmid also carried a CTG guide RNA (hereafter named gRNA#1) under the control of the constitutive *SNR52* promoter. The PAM used in this experiment was the TGG sequence located right at the border between the CTG tract and non-repeated DNA (Figure 1A). As controls, we used the same *Sp*Cas9-containing plasmid without the gRNA or a frameshift mutant of the *Sp*Cas9 gene resulting in a premature stop codon (*Sp*Cas9ΔNdeI) and the gRNA#1. The same genetic assay as previously was used (Mosbach et al., 2018; Richard et al., 2014). It is based on a modified suppressor tRNA gene (*SUP4*) in which a CTG trinucleotide repeat was integrated. The length of the CTG repeat at the start of the experiment was determined to be approximately 80 triplets. Four hours after transition from glucose to galactose medium, two faint bands were visible on a Southern blot, corresponding to the 5’ and 3’ ends of the *Sp*Cas9 DSB. No signal was detected in both control strains (Figure 1B). The DSB was quantified to be present in ca. 10% of the cells at any given time point and remained the same for the duration of the time course (4 hours). No evidence for repeat tract contraction was visible by Southern blot. Survival to the *Sp*Cas9 break was low (17.9%±4%), as calculated from CFU on galactose plates over CFU on glucose plates (Figure 2, see Materials & Methods). Surviving colonies were picked, total genomic DNA was extracted and the *SUP4*::CTG-locus was analyzed by Southern blot. Patterns observed were remarkably different among clones, most of them showing bands of aberrant molecular weight, either much larger or much shorter than the repeat tract. In some clones, a total absence of signal suggested that the probe target was deleted and in other cases weakness of the signal was compatible with partial deletion (Figure 3A). To understand these abnormal patterns, the genome of nine surviving clones were totally sequenced by paired-end Illumina. As a control, one clone in which *Sp*Cas9 had not been induced was also sequenced. In all nine cases, a deletion around the repeat tract was found, extending from a few nucleotides to several kilobases (Figure 3B). Some of the rearrangements involved flanking Ty1 retrotransposon LTRs, and in one case (clone #2), a complex event between a distant LTR (δ16) and the δ20 LTR close to the repeat tract was detected. Following this discovery, total genomic DNA was extracted from more surviving colonies, analyzed by Southern blot and rearrangement junctions were amplified by PCR and Sanger sequencing. Two different sets of primers were used to amplify the junction, su47/su48 allowing to amplify local rearrangements around the repeat tract, whereas su23/su42 were used to amplify larger rearrangements between Ty1 LTRs (Figure 4, Table 2). Surviving colonies were classified into 12 different types, according to the *SUP4*::CTG locus after *Sp*Cas9 induction: type I corresponded to a colony in which the repeat tract was unchanged, types II-V corresponded to local rearrangements around the repeat tract and types VI-XII corresponded to more extensive rearrangements, the last category involving a complex event between δ16 and δ20. A few examples of junction sequences are shown in Supplemental Figure 1.

**Figure 2:**
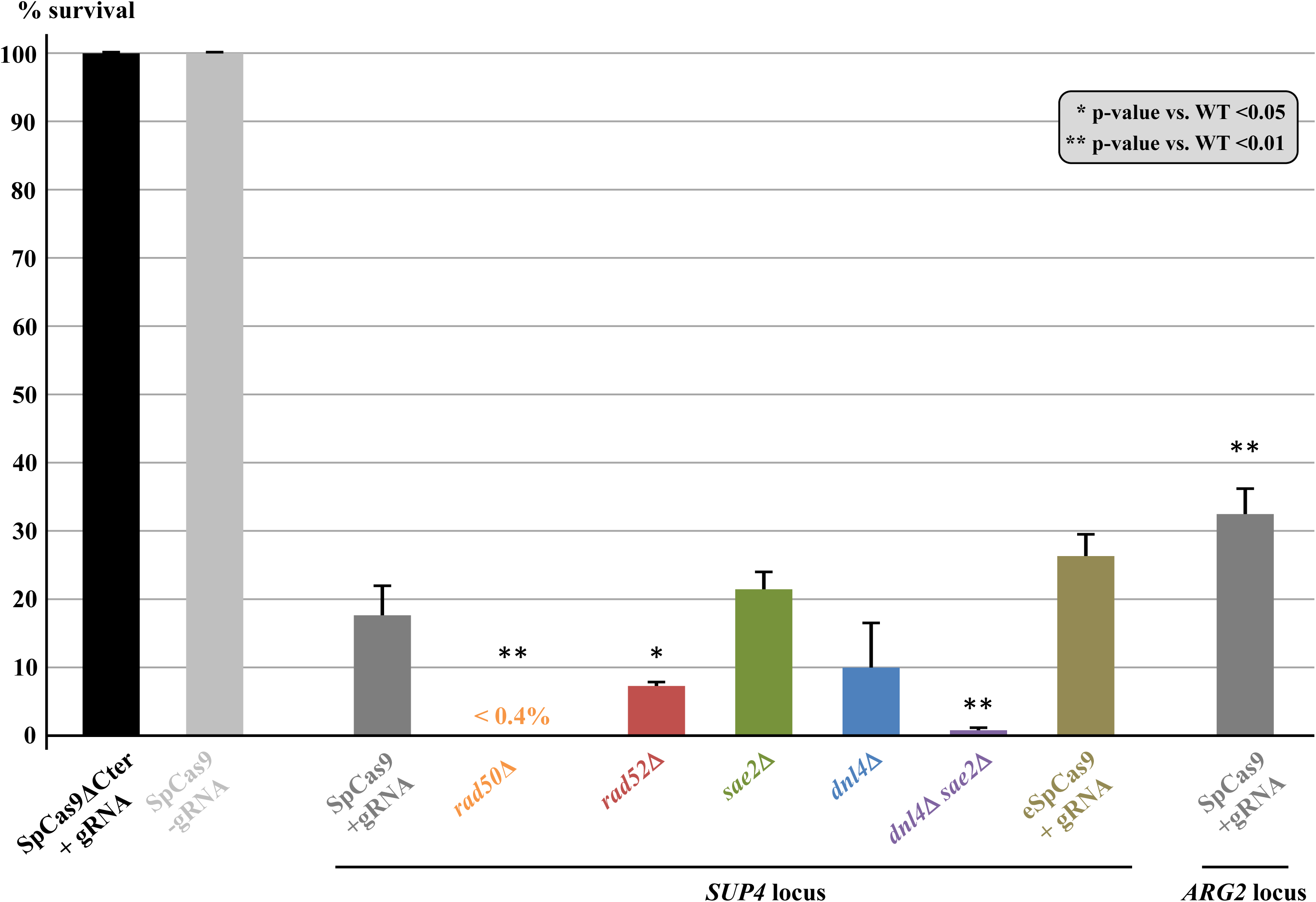
Yeast survival to Cas9 induction. For each strain, the same number of cells were plated on galactose and glucose plates and the survival was expressed in CFU number on galactose plates over CFU number on glucose plates. The mean and the 95% confidence interval are plotted for each strain. Significant t-test p-values when compared to wild-type *Sp*Cas9 survival are indicated by asterisks, as shown on the figure.

**Figure 3:**
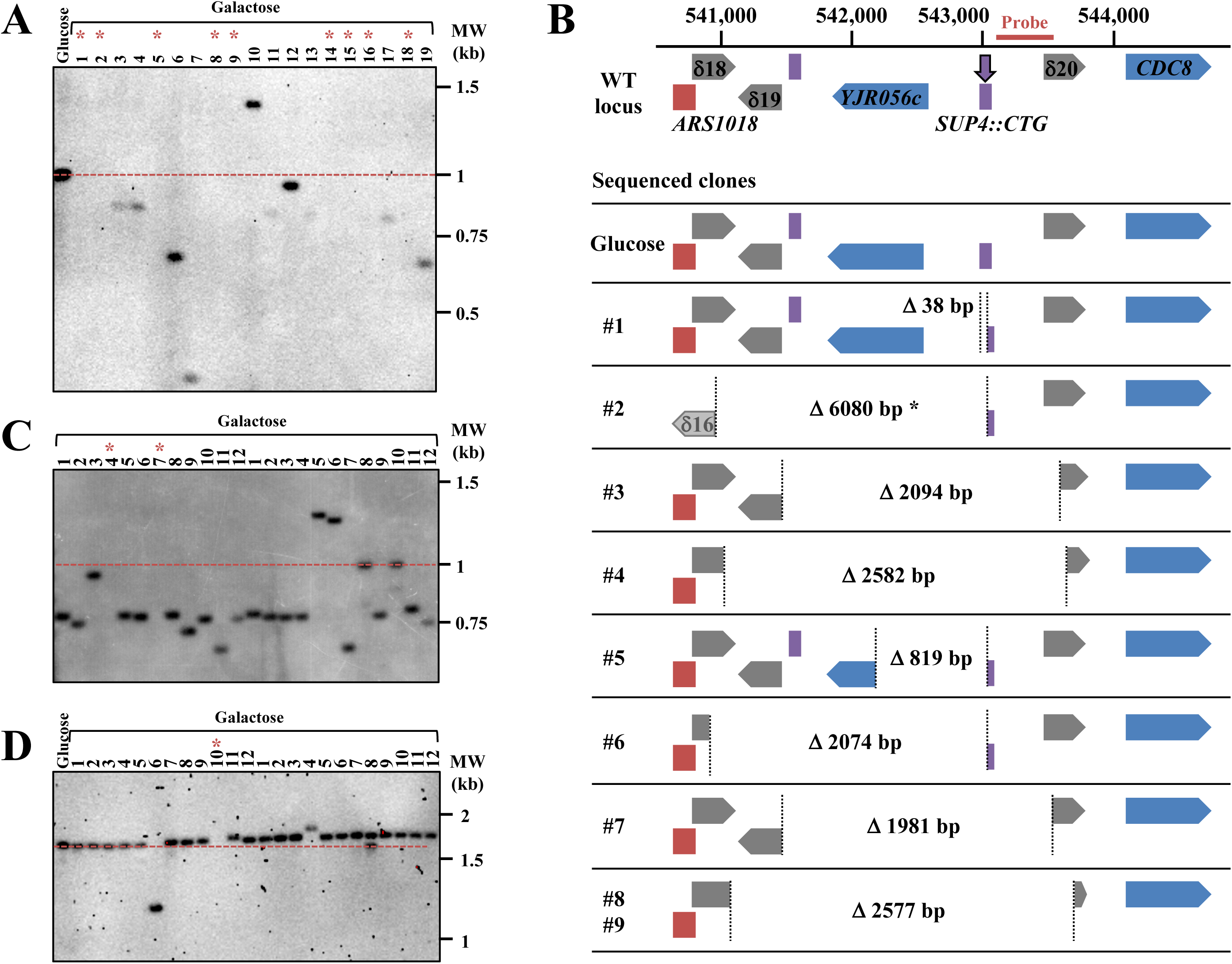
Chromosomal rearrangements following *Sp*Cas9 induction. **A**: Southern blot of genomic DNA at the *SUP4* locus in the wild-type strain. The probe hybridizes ∼300 bp downstream the repeat tract (see Figure 3B). The dotted red line shows the initial length of the CTG repeat tract. The lane labeled “Glucose” contains a clone in which Cas9 was not induced. Lanes numbered #1 through #19 contain independent clones in which Cas9 was induced. Asterisks point to lanes in which no signal was detected, meaning that the probe containing sequence was deleted. Note that signal intensities varies among lanes, showing that the probe did not fully bind to its target sequence, due to its partial deletion. **B**: Some examples of chromosome rearrangements following Cas9 induction in the wild-type strain. The genomic locus surrounding *SUP4* is shown on top, ARS1018 is drawn in red, delta elements are in grey, protein-coding genes are colored in blue and tRNA genes in purple. The DSB (vertical purple arrow) is induced within *SUP4*::(CTG)n. Chromosome coordinates are indicated above and the probe used for hybridization is represented by an horizontal red bar. The locus sequence was retrieved from the Saccharomyces Genome Database (http://yeastgenome.org/, genome version R64-2-1, released 18th November 2014). Under the reference locus are cartooned the different chromosomal structures observed in some of the survivors. A yeast colony that was grown in glucose was also sequenced as a control. For each clone, vertical dotted lines represent junctions of rearrangements observed, with deletion sizes indicated in base pairs. Asterisk: clone #2 showed a complex rearrangement with a local inverted duplication involving the δ16 LTR and the 3’ end of the *KCH1* gene 5 kb upstream *SUP4*. Two clones (#8 and #9) exhibit exactly the same chromosomal rearrangement at precisely the same nucleotides. Note that *CDC8* is an essential gene. **C**: Southern blot of genomic DNA at the *SUP4* locus in the *rad52*Δ strain. Legend as for Figure 3A. Note that for this Southern blot genomic DNA was digested with EcoRV (instead of Ssp I, see Methods), therefore the expected CTG repeat length was around 1.8 kb, instead of 1 kb. **D**: Southern blot of genomic DNA at the *SUP4* locus in the *sae2*Δ strain. Legend as for Figure 3A.

**Figure 4:**
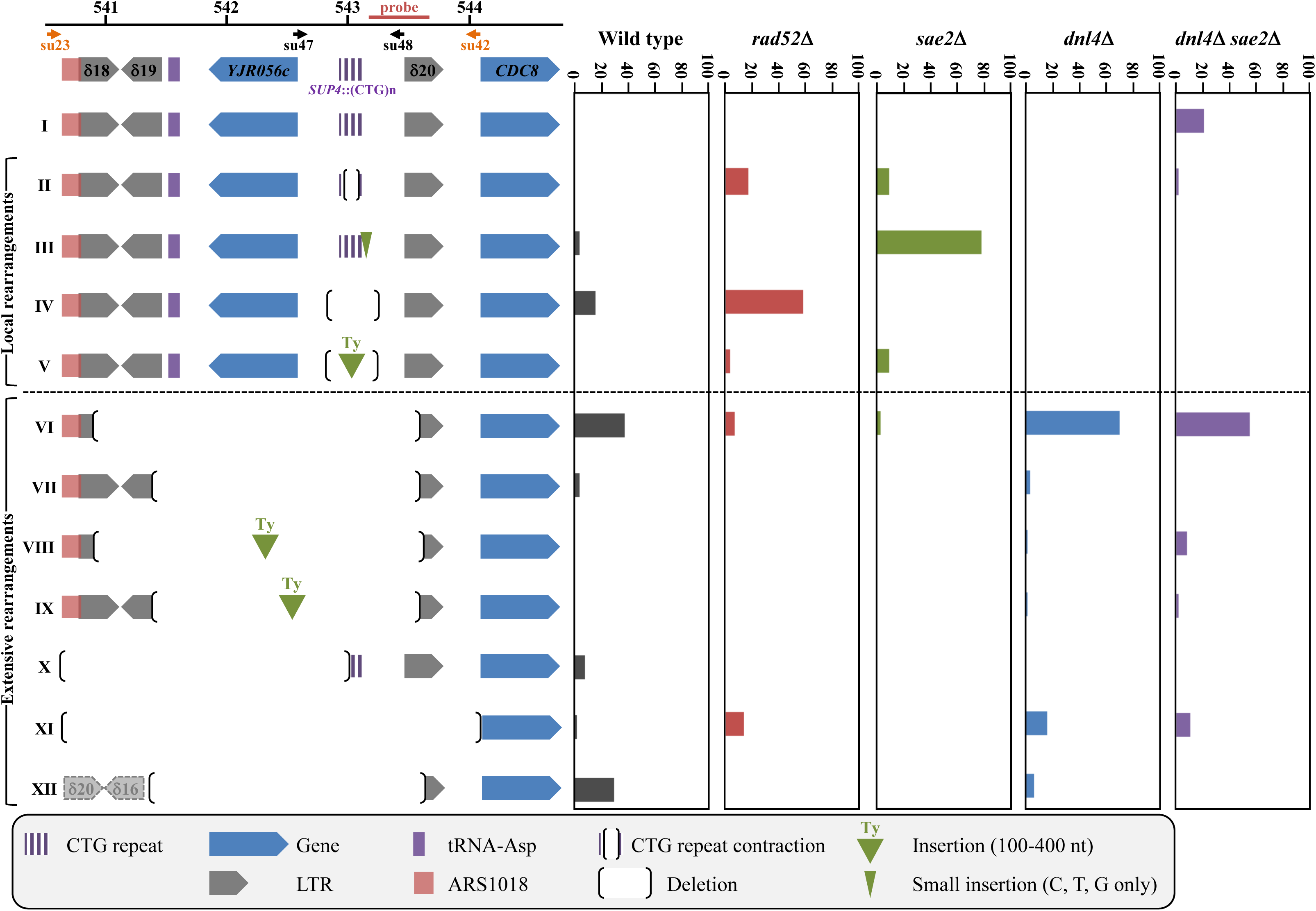
Summary of chromosomal rearrangements observed in wild-type and mutant strains, following *Sp*Cas9 induction. **Left**: The twelve different possible outcomes following *Sp*Cas9 induction are shown, subdivided in local and extensive rearrangements (see text for details). The *SUP4* locus is pictured and shows the position of each genetic element on yeast chromosome X. The probe used on Southern blots is shown, as well as both primer couples used to amplify the locus. In order to assess a given clone to a rearrangement type, the following rules were followed: i) when a band was detected by Southern blot, primers su47 and su48 were used to amplify the locus an sequence it. These events corresponded to types I-V. The absence of a PCR product indicated that primer su48 genomic sequence was probably deleted and therefore primers su23 and su42 were used to amplify and sequence the locus. These were classified as types IV-V events; ii) when no band was detected by Southern blot, primers su 23 and su42 were directly used to amplify and sequence the locus. These events were classified as types VI-X and XII. When no PCR product was obtained, it meant that at least one of the two primers genomic sequence was probably deleted and these events were classified as type XI. Note that this last category may also contain rare-but possible-chromosomal translocations that ended up in puting each primer in a separate chromosome, making unobtainable the PCR product. The extent of type XI deletions cannot go downstream the su42 primer, since the *CDC8* gene is essential. **Right**: The proportion of each type or event recovered is represented for wild type and mutants. Altogether, 220 surviving clones were sequenced, distributed as follows: WT: 51, *rad52*Δ: 29, *dnl4*Δ: 61, *sae2*Δ: 32, *dnl4*Δ *sae2*Δ: 47.

### Chromosomal rearrangements are under the control of RAD52, DNL4 and SAE2

We next decided to investigate the role of several genes known to be involved in DSB repair on chromosomal rearrangements generated by the *Sp*Cas9 nuclease. In a *rad52*Δ strain, in which all homologous recombination was abolished, survival significantly decreased below wild type (7.6%±0.7%, Figure 2). Molecular analysis of the survivors by Southern blot showed that rearrangements seemed to be less extensive than in the wild-type strain, fewer lanes showing a partial or total absence of signal (Figure 3C). Sequencing of the junctions confirmed that rearrangements between Ty LTRs were lost (Type VI events), except for two cases in which the deletion occurred through annealing of eight or nine nucleotides and was therefore *RAD52* independent (Figure 4 and Supplemental Figure 1, clones #C4 and #C7). This result shows that about 50% of colonies growing on galactose plates survived to the DSB by *RAD52*-dependent homologous recombination between two LTR elements flanking the trinucleotide repeat tract.

The possible role of non-homologous end-joining (NHEJ) in the observed rearrangements was also addressed by deleting the gene encoding yeast Ligase IV (*DNL4*). In the *dnl4*Δ strain, the level of detected DSBs was significantly lower than in wild type (Figures 1B and 1C), suggesting that DSB ends may be partially protected by the presence of Dnl4p, and were processed more rapidly when Ligase IV was absent. Survival was slightly decreased, but not significantly different from wild type (10.5%±6.3%, Figure 2). Molecular analysis of the survivors showed that local rearrangements were totally lost, whereas extensive rearrangements involving Ty LTR represented 84% of all events (Figure 4 and Supplemental Figure 1). Hence, we concluded that all local rearrangements were NHEJ dependent.

In a recent work, we showed that *SAE2* was essential to repair a DSB induced by a TALEN within a long trinucleotide repeat. In its absence, unrepaired breaks accumulated and DSB resection was lost on the trinucleotide repeat-containing end (Mosbach et al., 2018). We therefore tested the effect of a *sae2*Δ mutation on a *Sp*Cas9 DSB in the same experimental system. Southern blot analysis of repair intermediates showed that DSB ends accumulated twice as much in the *sae2*Δ mutant as compared to wild type (Figures 1B and 1C). In addition, a smear was detected below the 5’ DSB end (Figure 1B, orange bracket), hallmark of a resection defect (Chen et al., 2013). Survival was similar to wild type (21.5%±2.9%, Figure 2). Southern blot analysis of surviving colonies displayed very little size changes as compared to uninduced controls (Figure 3D). However, sequencing showed that the most frequent event was an insertion (or sometimes a small deletion) of one to eight nucleotides between the PAM and the repeat tract (Type III events, Figure 4). These local insertions represented 78% of all survivors, whereas only one Ty LTR recombination (Type VI) was detected (Supplemental Figure 1). This showed us that in the absence of *SAE2*, long range rearrangements were lost, probably due to the inability to resect the DSB into single-stranded DNA prone for homologous recombination.

The double mutant *sae2*Δ *dnl4*Δ was also built and showed an additive effect on survival, with a 30-fold reduction in CFU on galactose plates (0.6%±0.9%, Figure 2). This proved that in the absence of one of the two genes repair could occur by the other pathway, but absence of both genes was almost lethal to yeast cells receiving a *Sp*Cas9 DSB. Southern analysis showed that DSB levels were similar to *sae2*Δ levels (ca. 24% after 8 hrs versus 28% for *sae2*Δ), showing that *SAE2* was epistatic to *DNL4* for this phenotype. The smear corresponding to resection defects was also visible (Figure 1B, orange bracket). Interestingly, 21% of survivors exhibited zero to two triplets lost, which could be attributed to natural microsatellite instability. These were classified as Type I events and were specific of the *sae2*Δ *dnl4*Δ double mutant (Figure 4 and Supplemental Figure 1). It is possible that given the low survival rate, cells in which *Sp*Cas9 and/or the gRNA was mutated were positively selected during the time course in liquid culture and were therefore subsequently recovered on galactose plates. It is however surprising that such events were not recovered in *rad50*Δ cells. Remarkably, to the exception of the Type I events hereabove mentioned, all but one event corresponded to extensive rearrangements around the repeat tract, similarly to the single *dnl4*Δ mutant. This shows that although effects of both mutations were additive on cell survival, *DNL4* was epistatic to *SAE2*, when chromosomal rearrangements induced by *Sp*Cas9 were considered.

Finally, in a *rad50*Δ strain, the DSB accumulated over the duration of the time course at levels similar to *sae2*Δ mutants (Figures 1B and 1C). No smear was detected in this strain background, suggesting that the *sae2*Δ resection defect was specific of this gene and did not involve the integrity of the MRX-Sae2 complex. However, no survivor could be found on galactose plates, showing that the DSB was lethal in this mutant background (Figure 2).

In conclusion, when a *Sp*Cas9 DSB was induced into a long CTG trinucleotide repeat, cell survival was low and depended on *RAD50*, *RAD52*, *SAE2* and *DNL4*. Two classes of repair events were found: local rearrangements under the control of *DNL4* and therefore the NHEJ pathway, and extensive rearrangements under the control of *SAE2* and *RAD52*. In addition, the deletion of *RAD50* almost completely recapitulated the *sae2*Δ *dnl4*Δ double mutation, except that the smear was not visible in *rad50*Δ and no surviving cell could be found, suggesting that in the absence of this gene DSB repair cannot occur at all on CTG trinucleotide repeats, following a *Sp*Cas9-induced DSB.

### Enhanced SpCas9 generates the same chromosomal rearrangements as SpCas9

Over the last three years, several mutants of the widely used *Sp*Cas9 have been engineered or selected by genetic screens. *Sp*Cas9-HF1 and e*Sp*Cas9 were built to exhibit less off-target DSBs (Kleinstiver et al., 2016; Slaymaker et al., 2016), HypaCas9 was made to be even more accurate (Chen et al., 2017), Sniper-Cas9 also showed reduced off-target effects (Lee et al., 2018), while evoCas9 was selected in yeast for improved specificity (Casini et al., 2018). We decided to explore the possibility that chromosomal rearrangements observed in our experimental system were partly due to the fact that *Sp*Cas9 exhibited a high off-target activity on long CTG trinucleotide repeat tract, perhaps by generating more than one DSB within the repeat tract, or within the surrounding loci. In order to test this hypothesis, Enhanced *Sp*Cas9 (e*Sp*Cas9) was expressed in yeast, along with the same guide RNA as previously (gRNA #1, Figure 1A). Survival was slightly higher than with *Sp*Cas9 (26.3%±3.0%), but not significantly different (t test p-value= 0.06). DSB end accumulation was lower than *Sp*Cas9 (Figure 1B, 1C). Molecular analysis of surviving yeast cells did not show any statistical difference between types of rearrangements observed with e*Sp*Cas9 as compared to *Sp*Cas9 (Chi2 p-value= 0.14) (Figure 5 and Supplemental Figure 1). A second guide RNA (gRNA#2) was designed, so that the DSB would be made two nucleotides closer to the repeat tract end (Figure 1A). Interestingly, the number of rearrangements involving a LTR was lower than with gRNA#1 (19% with gRNA#2 vs 92% with gRNA#1) but the proportion of very large deletions (Type XI) significantly increased from 4% to 31% (Chi2 p-value= 1.6 10^-3^). We concluded that moving the DSB cut site two nucleotides toward non-repeated DNA increased the outcome of very large deletions. Altogether, these results show that using a more specific version of *Sp*Cas9 did not decrease chromosomal rearrangements, but actually increased extensive deletions, suggesting that the effects seen were probably not due to extra off-target DSBs within the CTG repeat tract or the surrounding loci.

**Figure 5:**
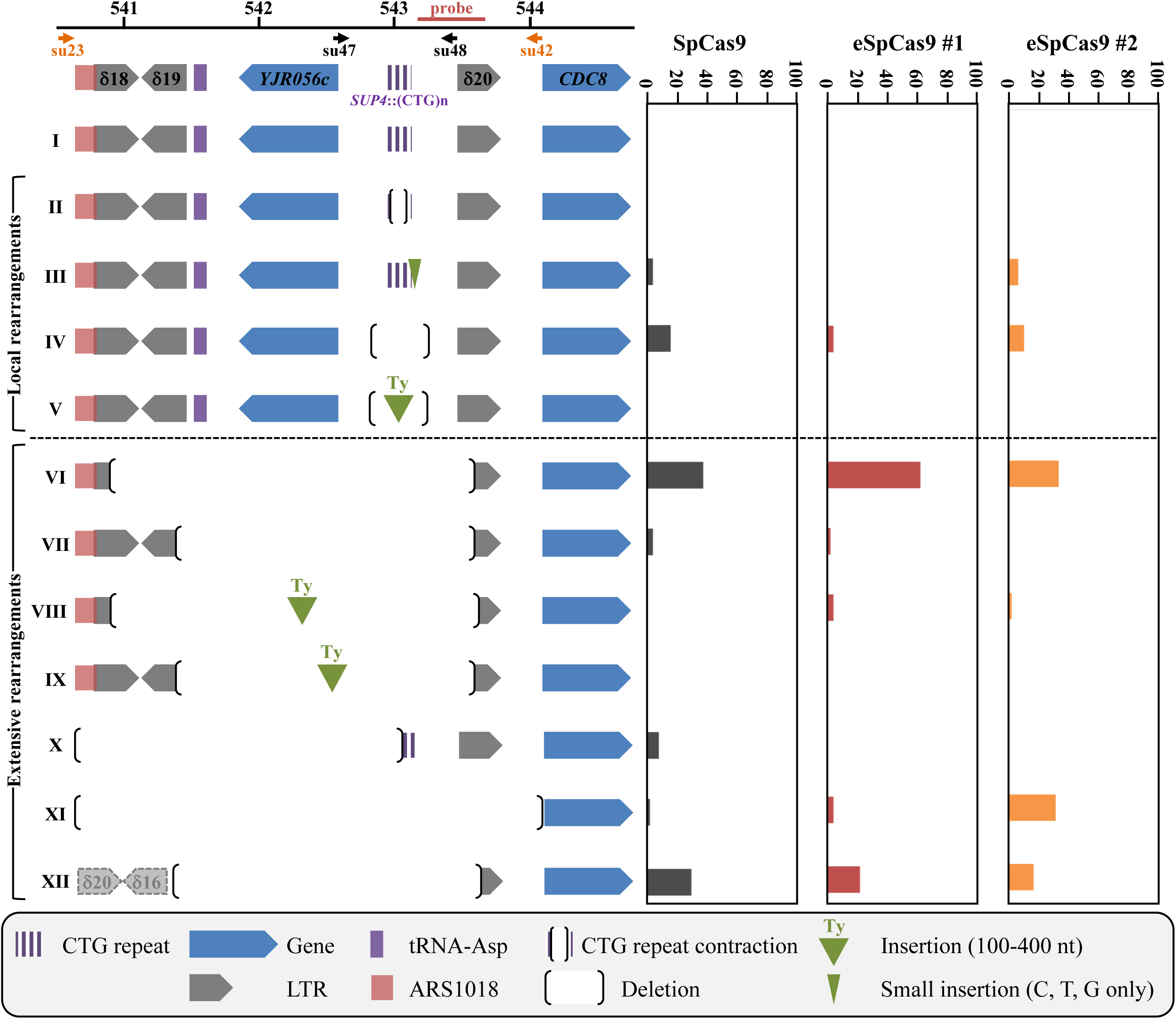
Summary of chromosomal rearrangements observed in following *Sp*Cas9 and enhanced *Sp*Cas9 (e*Sp*Cas9) inductions. See Figure 4 for legend. e*Sp*Cas9 #1 and #2 refer respectively to guide RNAs #1 and #2 described in Figure 1A. The following number of surviving clones were analyzed: *Sp*Cas9: 51, e*Sp*Cas9 #1: 49, e*Sp*Cas9 #2: 48. See text for details.

### Gene conversion efficacy is decreased when a Cas9 DSB is made within a long CTG trinucleotide repeat

Gene conversion is a very efficient DSB-repair mechanism in *S. cerevisiae*. We previously showed that a single DSB induced by the I-*Sce* I meganuclease in a yeast chromosome was efficiently repaired using a CTG repeat-containing homologous template as a donor (Richard et al., 1999a, 2000, 2003). In order to determine whether a Cas9-induced DSB within a CTG repeat was properly repaired by the recombination machinery, we reused a similar experimental system in which two copies of the *SUP4* allele were present on yeast chromosome X, one containing a (CTG)_60_ repeat tract and the other copy containing an I-*Sce* I recognition site (Figure 6A). In this ectopic gene conversion assay, 80.2%±2.3% of yeast cells survived to an I-*Sce* I DSB and 100% of survivors were repaired by gene conversion using the ectopic *SUP4*::(CTG)n copy as a donor (Richard et al., 2003). When *Sp*Cas9 was induced in the same yeast strain along with gRNA#1, only 32.6%±3.8% of CFU formed on galactose plates (Figure 2). Molecular analysis of surviving cells showed that 89% (34 out of 38) repaired by ectopic gene conversion, as expected, and now contain two I-*Sce* I recognition sites, one in each *SUP4* copy (Figure 6B, GC events). However, one expansion event was also detected, as well as one local rearrangement (Type IV) and two rearrangements involving a deletion and a DNA insertion (Type V). Intriguingly, the DNA insertion is a 211 bp piece of DNA from the *YAK1* gene, located 158 kilobases upstream the *ARG2* locus, on chromosome X left arm. This gene contains a long and imperfect CAG/CTG repeat within its reading frame, like many yeast genes (Field and Wills, 1998; Malpertuy et al., 2003; Richard and Dujon, 1996; Richard et al., 1999b). An unusual non-homologous recombination event occurred between the *YAK1* CAG/CTG repeat and the *ARG2* repeat, leading to a chimeric insertion (Figure 6C). This may be the result of an off-target DSB generated by *Sp*Cas9 within the *YAK1* repeat, or be due to an abnormal recombination event between the two CTG repeats following *Sp*Cas9 induction. Using the CRISPOR *in silico* tools, three off-target sites were found for *Sp*Cas9 gRNA#1 if no mismatch were allowed and 80 sites when up to three mismatches were permitted (Haeussler et al., 2016). The second best off-target score was *YAK1*. We concluded that gene conversion efficacy was reduced when the DSB was made by *Sp*Cas9 within a CTG repeat, as compared to an I-*Sce* I DSB made in a non-repeated sequence, partly due to occasional off-target DSBs in other CTG repeats of the yeast genome.

**Figure 6:**
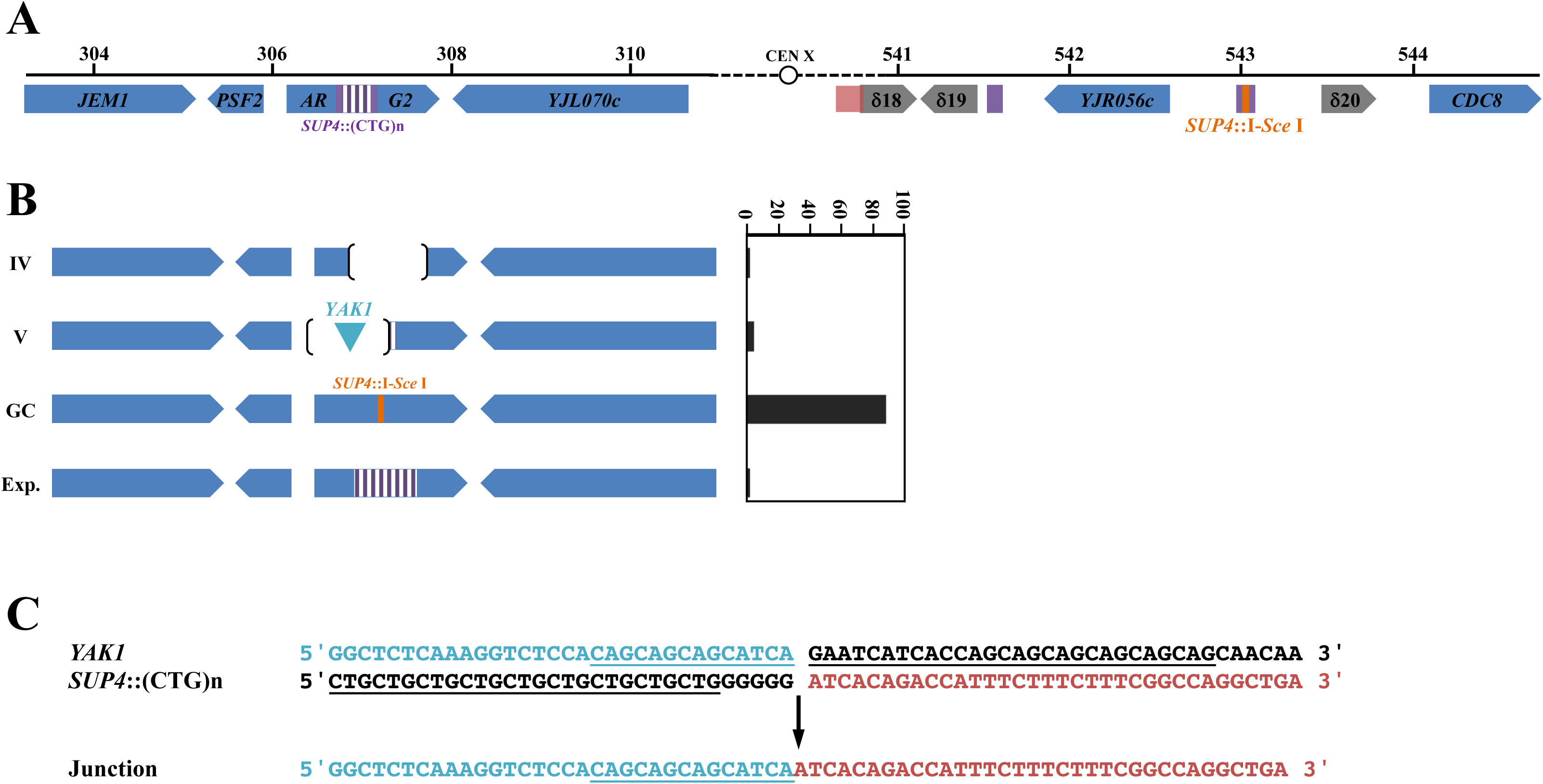
Chromosomal rearrangements observed at the *ARG2* locus, following *Sp*Cas9 induction. **A**: *ARG2* and *SUP4* loci drawn to scale. A 2.6 kb piece of DNA containing 1.8 kb of the *SUP4* locus in which a CTG repeat was integrated, as well as the *TRP1* selection marker were integrated at ARG2 (Richard et al., 2003). The *TRP1* gene is not represented here but is centromere-proximal located. **B**: Types of rearrangements observed. Types IV and V are explained in Figure 5. GC: gene conversion with *SUP4*::I-Sce I. Exp.: CTG repeat expansion. **C**: Type V rearrangements involving the *YAK1* gene. The imperfect repeat in *YAK1* and the CTG repeat in *SUP4* are underlined. The junction of the rearrangement contains the green sequence from *YAK1* ligated to the red sequence from *SUP4*::(CTG)n.

### Resection of a Cas9-induced double-strand break

Quantitative PCR experiments were performed in order to determine the resection level in strains in which Cas9 was induced. The nuclease generates a DSB in the very last CTG triplets of the repeat tract (Figure 1A). Therefore, the 5’ end of the break contains most of the 80 triplets whereas the 3’ end contains only two triplets. This allows to compare resection of a repeated and structured DNA end versus non-repeated DNA, concomitantly and in the same experimental setting. We took advantage of the convenient position of four *Eco*RV restriction sites, two on each side of the DSB, at different distances from the break (Figure 7A). Primers were designed in such a way that *Eco*RV digested DNA could not be PCR amplified. However, if DNA resection reached an *Eco*RV site, the resulting single-stranded DNA became resistant to digestion and therefore susceptible to amplification. In wild-type cells after eight hours, resection of the Cas9 DSB was always 100% at all *Eco*RV sites, except at the 3’ distal site in which it was a little lower, around 70% (Figure 7A). In *dnl4*Δ cells, resection was not statistically different from wild type. In the *rad50*Δ mutant, DSB resection was totally abolished on the 5’ end of the break that contains most of the repeat tract and severely impaired on the other end, showing that the MRX-Sae2 complex was essential in this process, on both DSB ends. Interestingly, the *sae2*Δ mutant exhibited a resection defect on the 5’ end of the break but not on the other side. This was also true for the double mutant *sae2*Δ *dnl4*Δ. All these data prove that: i) Ligase IV plays no role in DSB resection; ii) Sae2 is essential to resect a long CTG trinucleotide repeat but is dispensable to resect a non-repeated DSB end.

**Figure 7:**
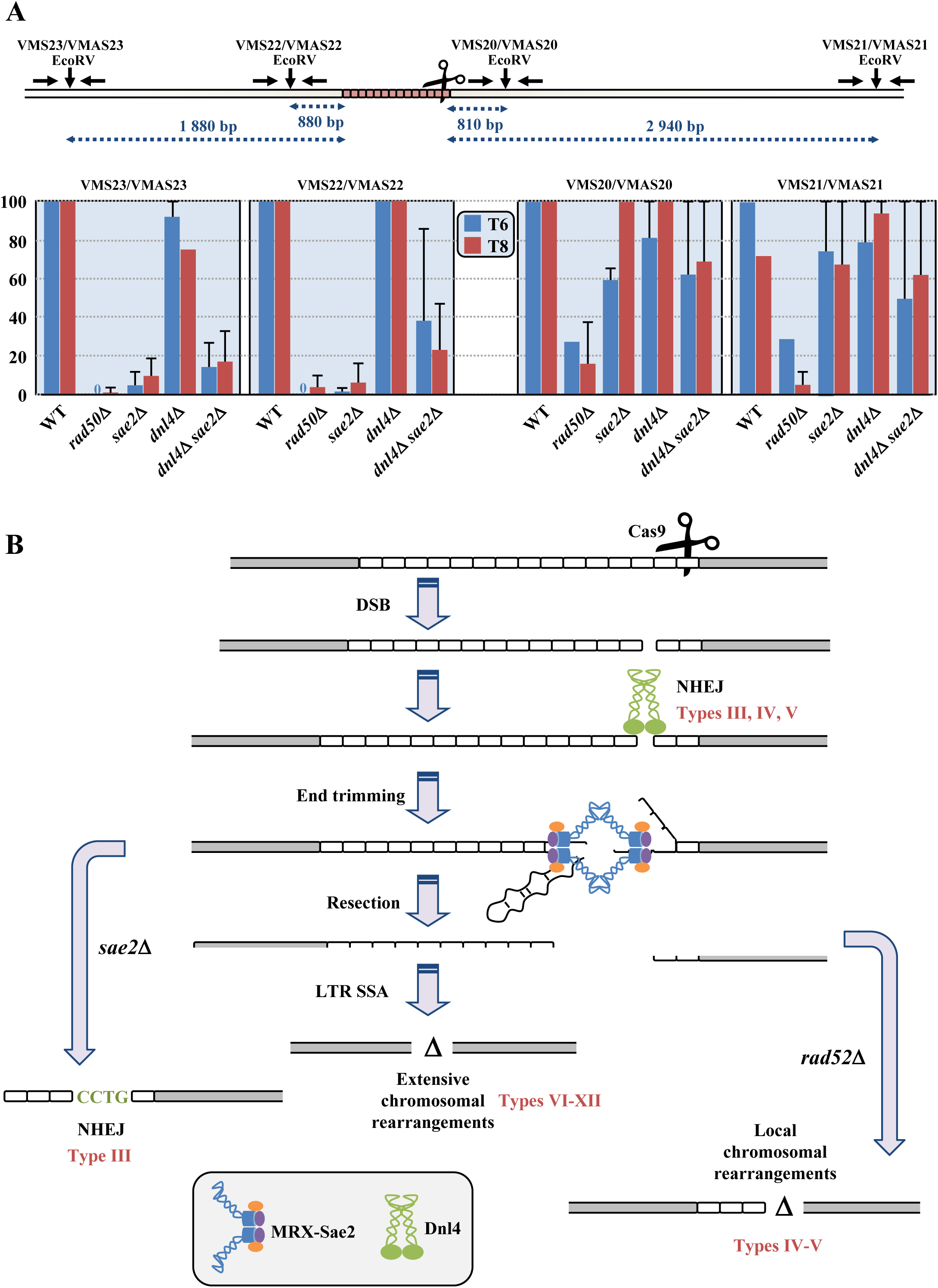
Quantification of double-strand break resection. **A**: Couples of primers used to amplify each EcoRV site are indicated above and can be found in Supplemental Table 2. EcoRV sites are shown by vertical arrows. Resection graphs are plotted for each primer pair. Average relative values of resection as compared to the total DSB amount detected on Southern blots are shown at 6 hours (in blue) and 8 hours (in red), along with standard deviations. **B**: Mechanistic model for chromosomal rearrangements following a Cas9-induced DSB. See text for details. Resulting rearrangement types are indicated in red near each pathway.

### Genome-wide mutation spectrum in cells expressing SpCas9

When carefully looking at deletion borders in haploid strains in which *Sp*Cas9 was induced, they were found to be more extensive on the 5’ side of the break than on the 3’ side (Figure 4). This suggests that larger 3’ deletions encompassing the essential gene *CDC8* or its promoter may not have been recovered because they would be lethal. This could also account for the lethality observed with *Sp*Cas9 in haploids. In order to check this hypothesis, we analyzed diploids containing *SUP4*::(CTG)n repeat tracts on both homologues, in which *Sp*Cas9 was induced. In these cells, both chromosomes could be cut by the nuclease. We quantified by qPCR *CDC8* copy number in six independent diploid survivors. In all cases, it was reduced by half as compared to a control qPCR on another chromosome (Supplemental Figure 2A). This showed that in these cells, only one of the two *CDC8* alleles was present, suggesting that the other was often deleted during DSB repair. In order to check if the whole chromosome could have been lost, we also amplified a region near the *JEM1* gene, on the other chromosomal arm, near the *ARG2* gene. Surviving clones showed a significantly higher signal, compatible with the presence of two chromosomes (Mann-Whitney-Wilcoxon rank test, p-value= 10^-3^). Therefore, it was concluded that the mortality observed in haploid cells expressing SpCas9 was at least partly due to the frequent deletion of the essential *CDC8* gene, but did not induce significant chromosome loss.

In order to detect possible off-target mutations, independent haploid and diploid colonies in which *Sp*Cas9 had been induced were deep-sequenced, using Illumina paired-end technology. In diploid cells, two nucleotide substitutions were detected out of five independent clones when *Sp*Cas9 was repressed (Supplemental Figure 2B). When the nuclease was induced, six mutations were detected out of 18 sequenced clones, a similar proportion. One 36-bp deletion was found in the *FLO11* minisatellite and two deletions of one repeat unit were found in AT dinucleotide repeats, but no mutation was found in any other CAG/CTG repeat tract. Altogether, we concluded that *Sp*Cas9 expression in diploid cells did not significantly increase genome-wide mutation frequency. The genome of 10 independent haploid cells in which *Sp*Cas9 was induced was also completely sequenced. Eight mutations were detected among six of these survivors, all of them being nucleotide substitutions in non-repeated DNA (Supplemental Figure 2B). This is statistically not different from what was observed in diploids (Fisher exact test p-value= 0.003). We concluded that besides chrosomosomal rearrangements around the *SUP4* locus observed in these haploids, *Sp*Cas9 did not induce other mutations in yeast cells in which it was expressed.

## Discussion

### SpCas9-induced DSB repair within CTG repeats generates chromosomal rearrangements

In previous works, in which we induced a DSB within a CTG repeat using a dedicated TALEN, 100% of surviving yeast colonies repaired the break by contracting the repeat tract (Richard et al., 2014). These contractions occurred by an iterative single-strand annealing mechanism, that depended on *RAD52*, *RAD50* and *SAE2*, but was independent of *LIG4*, *POL32* and *RAD51* (Mosbach et al., 2018). It was therefore striking and completely unexpected that a DSB made by the *Sp*Cas9 nuclease (or by its more specific mutant version, e*Sp*Cas9) at exactly the same location within the very same CTG trinucleotide repeat induced frequent chromosomal rearrangements around the repeat tract and almost no repeat contraction. With the TALEN, repeat contraction was shown to be an iterative phenomenon, involving several rounds of cutting and contraction until the repeat tract was too short for the two TALEN arms to dimerize and induce a DSB (Mosbach et al., 2018; Richard et al., 2014). A similar outcome was expected with *Sp*Cas9, iterative rounds of cutting and contraction could occur until the remaining CTG repeat tract would be too short for the gRNA to bind and induce a DSB. However, this was not observed here, surprisingly proving that a *Sp*Cas9-induced DSB was differently repaired from a TALEN-induced DSB targeting the same exact repeated sequence.

Spontaneous homologous recombination events between delta elements surrounding *SUP4* were already described by the past (Rothstein et al., 1987). However, in the present case, Cas9-induced deletions also involved microhomology sequences or no homology at all, in addition to delta LTR elements, suggesting that the initiating damage was different in both experimental systems (replication-induced single-strand nicks vs. nuclease-induced double-strand breaks, for example). Our data are more reminiscent of a previous work in which spontaneous deletions around the *URA2* gene were classified in seven different classes, six of them harboring microhomologies at their junctions and one showing no obvious homology (Welcker et al., 2000). In a recent work, using a GFP reporter system in human cells, it was shown that SpCas9 induced contractions as well as expansions of long CTG trinucleotide repeats, whereas the nickase mutant SpCas9-D10A only induced contractions. However, it was not possible to determine whether local rearrangements could be present in some of the surviving cells (Cinesi et al., 2016). In a recent work looking at the effect of a Cas9-induced DSB at the *LYS2* locus in *S. cerevisiae*, the authors found frequent *POL4*-dependent small insertions (1-3 bp) in 42-68% of the survivors (depending on the PAM used) and local deletions (1-17 bp) in the remaining cases. In the present experiments, local rearrangements account for 19.6% with 20% of those being small insertions (Figure 4). The remaining events (80.4%) corresponded to extensive rearrangements involving retrotransposon LTRs. However, given that there is no transposon or transposon remnant in the close proximity of the *LYS2* locus, the authors could not retrieve LTR rearrangements (Lemos et al., 2018). This strongly suggests that rearrangements observed heavily depend on the surrounding chromosomal location where the DSB is made.

When an I-*Sce* I DSB was induced within a *SUP4* allele, the break could be repaired by gene conversion with a CTG repeat-containing homologous donor at the *ARG2* locus. All yeast cells repaired by gene conversion with the donor, generating repeat contractions and expansions in the process (Richard et al., 2003). Here, the exact reverse reaction was induced, the break was made within CTG repeats and repaired with a non-repeated sequence. DSB repair was much less efficient, since only 32.6% of the cells survived (Figure 2) and less specific since 10% of the repair events were unfaithful recombination (Figure 6B). This shows that when a Cas9 DSB was made into a CTG repeat, gene conversion was impaired, either by the repeat tract or by the Cas9 protein, or by both.

### Ligase IV and Sae2 are respectively driving local and extensive chromosomal rearrangements

Yeast Ligase IV is encoded by the *DNL4* gene and is the enzyme used to ligate DSB ends during non homologous end-joining (Wilson et al., 1997). It was previously shown that *RAD50* and *SAE2* were essential to resect and process a TALEN-induced DSB but a *DNL4* deletion had no effect on break processing, cell survival or repair efficacy (Mosbach et al., 2018). On the contrary, repair of a Cas9, an HO or an I-*Sce* I DSB at the *MAT* locus, in the absence of any homologous donor casette, was shown to be dependent on the product of the *DNL4* gene (Frank-Vaillant and Marcand, 2001; Lemos et al., 2018). *Sp*Cas9 DSB repair has also been studied in human cells in the presence of a drug (NU7441), acting as a chemical inhibitor of non-homologous end-joining. In these conditions, the frequency of single-base insertions and small deletions decreased whereas larger deletions increased, suggesting that these repair events occurred by an alternative end-joining mechanism (alt-EJ/MMEJ) involving microhomologies flanking the DSB (Charpentier et al., 2018; van Overbeek et al., 2016). Here, we showed that when *DNL4* was inactivated, local deletions were totally lost. However, survival was not significantly decreased because yeast cells could repair the DSB using LTR recombination, generating extensive deletions around the repeat tract (Figure 4). Supporting this model, the absence of any resection defect in the *dnl4*Δ mutant proved that in the absence of end-joining, resection may take place very efficiently to repair the DSB by homologous recombination, using flanking homologies.

*SAE2* is associated to the *MRE11*-*RAD50*-*XRS2* complex, whose roles are multiple during DSB repair (Haber, 1998) and it was proposed to encode an endonuclease activity essential to process DNA hairpins (Lengsfeld et al., 2007), as well as to resect I-*Sce* I double-strand breaks (Mimitou and Symington, 2008). We previously showed that it was essential to resect a TALEN-induced DSB end containing a long CTG trinucleotide repeat, but less important to resect non-repeated DNA (Mosbach et al., 2018). In the present experiments, extensive rearrangements involving LTR elements were lost in a *sae2*Δ mutant, and 97% of yeast cells repaired the DSB by local rearrangements, most of them resulting in insertions or deletions between the PAM and the gRNA sequence (Figure 4), inactivating *Sp*Cas9 capacity to induce another DSB. Small insertions of a few nucleotides were also frequently detected following *Sp*Cas9 DSB induction at the VDJ locus in human B cells (So and Martin, 2019) or at the *MAT* locus in *S. cerevisiae* (Lemos et al., 2018). However, in our experiments, all nucleotides inserted were C, T or G, all three encoded by the gRNA. No insertion of an adenosine residue was found out of 28 insertions sequenced (Supplemental Figure 1). This intriguing observation suggests the possibility that the gRNA could be used as a template to repair the DSB, as it was demonstrated that a single-stranded RNA could be used to repair an HO-induced DSB into the *LEU2* gene (Storici et al., 2007).

Although *DNL4* and *SAE2* trigger different types of chromosomal rearrangements and none of the single mutants significantly decreased survival, the *dnl4*Δ *sae2*Δ double mutant abolished repair, almost to the point of the *rad50*Δ mutant, since only 0.6% of the cells survived (Figure 2), showing the synthetic effect of both mutations. However, repair events in the double mutant were similar to those observed in *dnl4*Δ, proving that *DNL4* was epistatic on *SAE2* (Figure 4). All these results are compatible with a model in which a Cas9 DSB was tentatively repaired by NHEJ first (Figure 7B). Then, if repair was unsuccessful or if *DNL4* was inactivated, resection proceeded with MRX-Sae2 in charge of removing secondary structures present at DSB ends. When resection reached flanking LTRs, repair occured by *RAD52*-mediated SSA. In the absence of this gene, the break was repaired by *RAD52*-independent local rearrangements. Finally, when *SAE2* was inactivated, resection was impeded on the 5’ DSB end containing most of the CTG repeat tract (Figure 7A), and mutagenic NHEJ was favored, leading to local insertions and deletions. It is unknown whether Sae2 would play the same essential role on other secondary structure-forming trinucleotide repeats, like GAA or CGG triplets, or if its activity is specific of CTG triplets, hence of a structure rather than a repeat. It is also unclear whether other nucleases, like *EXO1* or *DNA2*, would be important to perform long range resection on a long CTG repeat tract (Mimitou and Symington, 2008; Zhu et al., 2008), but the present experimental system allows us to address this question in a unique and unbiased way.

## Methods

### Yeast strains and plasmids

All mutant strains were built from strain GY6162-3D by classical gene replacement method (Orr-Weaver et al., 1981), using KANMX4 or *HIS3* as marker (Supplemental Table 1). KANMX4 cassettes were amplified from the EUROSCARF deletion library, using primers located 1kb upstream and downstream the cassette. VMS1/VMAS1 were used to amplify *rad52*Δ::KANMX, VMS2/VMAS2 were used to amplify *rad51*Δ::KANMX, VMS3/VMAS3 were used to amplify *pol32*Δ::KANMX, VMS4/VMAS4 were used to amplify *dnl4*Δ::KANMX, VMS6/VMAS6 were used to amplify *rad50*Δ::KANMX and SAE2up/SAE2down were used to amplify *sae2*Δ::KANMX (Supplemental Table 2). VMY350 and VMY352 strains were respectively used to construct VMY650 and VMY352 by mating-type switching, as follows: the pJH132 vector (Holmes and Haber, 1999) carrying the HO endonuclease under the control of an inducible *GAL1-10* promoter was transformed in the haploid *MAT*α strains. After 5h of growth in lactate medium, HO expression was induced by addition of 2% galactose (final concentration) and grown for 1.5 hour. Cells were then plated on YPD and mating type was checked three days later by crosses with both *MAT*a and *MAT*α tester strains.

For *Sp*Cas9 inductions, addgene plasmid #43804 containing the nuclease under the control of the GalL promoter and the *LEU2* selection marker was digested with HpaI and cloned into yeast by homology-driven recombination (Muller et al., 2012) with a single PCR amplified fragment containing the *SNR52* promoter, the gRNA#1 and the *SUP4* terminator, using primers SNR52Left and SNR52Right (Supplemental Table 2) to give plasmid pTRi203. A frameshift was then introduced in this plasmid by NdeI digestion followed by T4 DNA polymerase treatment and religation of the plasmid on itself, to give plasmid pTRi206. In this plasmid, the *Sp*Cas9 gene is interrupted by a stop codon after amino acid Ile_161_. The haploid GFY6162-3D strain (or its mutant derivatives), was subsequently transformed with pTRi203 or pTRi206 and transformants were selected on SC-Leu. The plasmid containing Enhanced *Sp*Cas9 (version 1.1, Addgene #71814, Slaymaker et al., 2016) was a generous gift of Carine Giovannangeli from the *Museum National d’Histoire Naturelle*. The e*Sp*Cas9 gene was amplified using primers LP400 and LP401 (Supplemental Table 2) and cloned into yeast cells in the Addgene#43804 plasmid digested with BamHI, by homology-driven recombination, with 34-bp homology on one side and 40-bp homology on the other side (Muller et al., 2012), to give plasmid pLPX11. For the gRNA#1, plasmid pLPX11 was digested with HpaI and cloned into yeast by homology-driven recombination (Muller et al., 2012) with a single PCR amplified fragment containing the *SNR52* promoter, the gRNA#1 and the *SUP4* terminator, as above to give plasmid pTRi207. For the gRNA#2, a guide RNA cassette was ordered from ThermoFisher (GeneArt), flanked by *Eco*RI sites and was cloned in pRS416 (Sikorski and Hieter, 1989) using standard procedures to give plasmid pLPX210.

### In silico simulations of off-target sites

To assess the number of off-target sites for *Sp*Cas9 in *Saccharomyces cerevisiae*, online tools were used. CRISPOR is a software that evaluates the specificity of a guide RNA through an alignment algorithm that maps sequences to a reference genome to identify putative on- and off-target sites (Li and Durbin, 2009). To predict off-target sites, the online tool sequentially introduces changes in the sequence of the gRNA and checks for homologies in the specified genome (Haeussler et al., 2016).

### Cas9 inductions

Before nuclease induction, Southern blot analyses were conducted on several independent subclones to select one containing ca. 80 CTG triplets. For Cas9 inductions, yeast cells were grown overnight at 30°C in liquid SC-Leu medium, then washed with sterile water to remove any trace of glucose. Cells were split in two cultures, half of the cells were grown in synthetic-Leu medium supplemented with 2% galactose (final concentration) and the other half were grown in synthetic-Leu medium supplemented with 2% glucose (final concentration). Around 4×10^8^ cells were collected at different time points (T=0, 4, 5, 6, 7 and 8 hours) and killed by addition of sodium azide (0.01% final). Cells were washed with water, and frozen in dry ice before DNA extraction. To determine survival to Cas9 induction, 24 hours after the T0 time point, cells were diluted to an appropriate concentration, then plated on SC-Leu plates containing either 20 g/l glucose or galactose. After 3-5 days of growth at 30°C, ratio of CFU on galactose plates over CFU on glucose plates was considered to be the survival rate.

### Double-strand break analysis and quantification

Total genomic DNA (4 μg) of cells collected at each time point was digested for 6h with EcoRV (40 U) (NEB) loaded on a 1% agarose gel (15×20 cm) and run overnight at 1 V/cm. The gel was vaccum transfered in alkaline conditions to a Hybond-XL nylon membrane (GE Healthcare) and hybridized with two randomly-labeled probes specific of each side of the repeat tract, upstream and downstream the *SUP4* gene (Viterbo et al., 2018). After washing, the membrane was overnight exposed to a phosphor screen and signals were read and quantified on a FujiFilm FLA-9000.

### SUP4 locus analysis after Cas9 induction

Several colonies from each induced or repressed plates were picked, total genomic DNA (4 μg) was extracted with Zymolyase, digested for 6h by SspI (20 U) (NEB), loaded on a 1% agarose gel (15×20 cm) and run overnight at 1V/cm. The gel was vaccum transfered in alkaline conditions to a Hybond-XL nylon membrane (GE Healthcare) and hybridized with a randomly-labeled PCR fragment specific of a region downstream the *SUP4* gene, amplified from the su8-su9 primer couple (Supplemental Table 2). After washing, the membrane was overnight exposed on a phosphor screen and signals were revealed on a FujiFilm FLA-9000. Genomic DNA of each clone for which a signal was detected by Southern blot was subsequently amplified with su47-su48 primers and sequenced using su47 (Supplemental Table 2). Genomic DNA of clones for which no signal was detected by Southern blot of no PCR product was obtained with su47-su48 were subsequently amplified with su23-su42 primers and sequenced using su42 (Supplemental Table 2). Sanger sequencing was performed by GATC biotech.

### Analysis of Cas9-induced DSB end resection by qPCR

A real-time PCR assay, using primer pairs flanking EcoRV sites 0.81 kb and 2.94 kb away from the 3’ end of the CTG repeat tract (VMS20/VMAS20 and VMS21/VMAS21 respectively) and 0.88 kb and 1.88 kb away from the 5’end of the CTG repeat tract (VMS22/VMAS22 and VMS23/VMAS23 respectively), was used to quantify end resection. Another pair of primers was used to amplify a region of chromosome X near the *ARG2* gene (Viterbo et al., 2016), to serve as an internal control of DNA amount (JEM1f-JEM1r). Genomic DNA of cells collected at T=0h, T=6h and T=8h was split in two fractions, incubated at 80°C for 10 minutes in order to inactivate any remaining active DNA nuclease, then one fraction was used for EcoRV digestion and the other one for a mock digestion in a final volume of 15 μl. Samples were incubated for 5h at 37°C, then the enzyme was inactivated for 20 min at 80°C. DNA was subsequentltly diluted by adding 55 μl of ice-cold water, and 4 μ was used for each real-time PCR reaction in a final volume of 25 μl. PCRs were performed with the Absolute SYBR Green Fluorescein mix (Thermo Scientific) in the Mastercycler S realplex (Eppendorf), using the following program: 95°C 15min, 95°C 15sec, 55°C 30 sec, 72°C 30 sec repeated 40 times, followed by a 20 min melting curve. Reactions were performed in triplicates and the mean value was used to determine the amount of resected DNA, using the following formula: raw resection=2/(1+2^ΔCt^) with ΔCt=C_t,EcoRV_-C_t,mock_. Relative resection values were calculated by dividing raw resection values by the percentage of DSB quantified at the corresponding time point.

The same protocol was used to determine the relative amount of *CDC8* and chromosome X in surviving clones after Cas9 induction, except that total genomic DNA was not digested prior to real-time PCR. Primer couples VMS23-VMAS23 were used to amplify *CDC8* and JEM1f-JEM1r for chromosome X left arm. Primers Chromo4_f and Chromo4_r were were used to amplify a region of chromosome IV as an internal control for total DNA amount. See Supplemental Table 2 for all primer sequences.

### Library preparation for deep-sequencing

Approximately 10 μg of total genomic DNA was extracted and sonicated to an average size of 500 bp, on a Covaris S220 (LGC Genomics) in microtubes AFA (6×16 mm) using the following setup: Peak Incident Power: 105 Watts, Duty Factor: 5%, 200 cycles, 80 seconds. DNA ends were subsequently repaired with T4 DNA polymerase (15 units, NEBiolabs) and Klenow DNA polymerase (5 units, NEBiolabs) and phosphorylated with T4 DNA kinase (50 units, NEBiolabs). Repaired DNA was purified on two MinElute columns (Qiagen) and eluted in 16 μl (32 μl final for each library). Addition of a 3’ dATP was performed with Klenow DNA polymerase (exo-) (15 units, NEBiolabs). Home-made adapters containing a 4-bp unique tag used for multiplexing, were ligated with 2 μl T4 DNA ligase (NEBiolabs, 400,000 units/ml). DNA was size fractionated on 1% agarose gels and 500-750 bp DNA fragments were gel extracted with the Qiaquick gel extraction kit (Qiagen). DNA was PCR amplified for 12 cycles with Illumina primers PE1.0 and PE2.0 and Phusion DNA polymerase (1 unit, Thermo Scientific). Six PCR reactions were pooled for each library, and purified on a Qiagen purification column. Elution was performed in 30 μl and DNA was quantified on a spectrophotometer and on agarose gel.

### Analysis of paired-end Illumina reads

Multiplexed libraries were loaded on a HiSeq2500 (Illumina), 110 bp paired-end reads for haploids and 260 bp paired-end reads for diploids were generated. Reads quality was evaluated by FastQC v.0.10.1 (http://www.bioinformatics.babraham.ac.uk/projects/fastqc/). Reads were mapped along S288C chromosome reference sequence (*Saccharomyces Genome Database,* release R64-2-1, November 2014), using the paired-end mapping mode of BWA v0.7.4-r385 with default parameters (Li and Durbin, 2009). The output SAM files were converted and sorted to BAM files using SAMtools v0.1.19-44428cd (Li et al., 2009). The command *IndelRealigner* from GATK v2.4-9 (DePristo et al., 2011) was used to realign the reads. Duplicated reads were removed using the option “*MarkDuplicates”* implemented in Picard v1.94 (http://picard.sourceforge.net/). Reads uniquely mapped to the reference sequence with a minimum mapping quality of 20 (Phred-scaled) were kept. Mpileup files were generated by SAMtools without BAQ adjustments. SNPs and INDELs were called by the options “*mpileup2snp*” and “*mpileup2indel*” of *Varscan2 v*2.3.6 (Koboldt et al., 2012) with a minimum depth of 10 reads for haploids and 20 reads for diploids. Average read coverage was 255X for diploid cells (σ= 187X) and 190X for haploids (σ= 43X). Diploid strains are homozygous except for selection markers and some specific loci like *MAT* and *SUP4*. Therefore, *de novo* heterozygous mutations should represent 50% of reads, on the average. Taking that into account, lower and upper thresholds for variant allele frequency were respectively set between 30% and 70% in diploids. For haploids, the threshold for minimum variant allele frequency was set at 70%. Mutations less than 10 bp away from each other were discarded to avoid mapping problems due to paralogous genes or repeated sequences. To assess microsatellite mutations, we only retained reads uniquely anchored at least 20 bp on each side of the microsatellite (Fungtammasan et al., 2015). All detected mutations were manually examined using the IGV software (version 2.3.77), and compared between all sequenced libraries for interpretation. All the scripts used in order to process data are available on github (https://github.com/sdeclere/nuclease). All Illumina sequences were uploaded in the European Nucleotide Archive (ENA), accession number PRJEB16068.

## Ackowledgments

We thank Carine Giovannangeli for the generous gift of the Enhanced *Sp*Cas9 plasmid. V. M. was supported by Fondation Guy Nicolas and Fondation Hardy. L. P. was the recipient of a graduate student CIFRE fellowship from SANOFI. W. V.-Z. is the recipient of a PhD fellowship from la Ligue Nationale Contre le Cancer. This work was generously supported by the Institut Pasteur and by the Centre National de la Recherche Scientifique (CNRS).

## Author contributions

V. M. built yeast mutant strains, performed time courses, Southern blots, survival experiments, Illumina library constructions and the initial resection assays. D. V. built yeast mutant strains, performed time courses, Southern blots and survival experiments. L. P. built pLPX11 and pLPX210 plasmids and did the first *Sp*Cas9 induction as well as survival experiments. W. V.-Z. performed resection assays. S. D. D. analyzed Illumina sequences. G.-F. R. designed the experiments, analyzed rearrangement junction sequences and wrote the manuscript.

---

**Supplemental Figure 1:** Sequences at the left and right of junctions in rearranged haploid clones. Junctions were deduced from Illumina read mapping (when available) and confirmed by subsequent PCR and Sanger sequencing. Nucleotides in red are those used to anneal each DSB end, and are therefore present in only one copy in the genomic sequence. The extent of calculated deletions (Δ) is indicated in parentheses. Nucleotides in red in parentheses correspond to small deletions. Nucleotides in green correspond to insertions. The length of Ty insertions is indicated along with the LTR it comes from. Nucleotides in purple (*Sp*Cas9 at the *ARG2* locus) correspond to the I-*Sce* I site (see text). Nucleotides in light blue correspond to homeologies between the left and right junction sequences that were lost after rearrangement (the junction sequence shows the nucleotide in blue, not the one in red). Nucleotides in light blue in parentheses correspond to homeologies that were deleted during the rearrangement. Note that extended homologies between LTRs does not always allow to determine the exact breakpoint with a high precision.

**Supplemental Figure 2: Genome-wide mutation spectrum observed in haploid and diploid cells following Cas9 induction**

**A**: Real-time PCR quantification of *CDC8* and *JEM1* amounts relative to an internal control on chromosome IV, in diploid cells in which Cas9 was induced. Half the amount of *CDC8* product was detected in each clone analyzed. This was significantly different from the amount of product amplified from the *JEM1* gene located on the other chromosome X arm.

**B**: Illumina results for diploid and haploid cells. For each clone, the number of mutations detected is shown. Substit.: nucleotide substitution; Indel: insertion or deletion; Indel micro.: insertion or deletion of one repeat unit in a microsatellite. The asterisk corresponds to a 36 bp deletion in the *FLO11* minisatellite (36 bp repeat).

