## Supplemental Figure 1 for "Resection and repair of a Cas9 double-strand break at CTG trinucleotide repeats induces local and extensive chromosomal rearrangements"

**Wild-type *Sp*Cas9 junctions (at the *SUP4* locus)**

| | |
|---|---|
| Parental allele | GGAATG CTG$_{90}$ CTGGGGGGATCACAGACCAT |
| #2 | GGAATG CTG$_{21}$ (A)TGCTGGGGGGATCACAGACCAT |
| #23 | GGAATG CTG$_{82}$ (C)TGCTGGGGGGATCACAGACCAT |
| #11 | GGAATG CTG$_{5}$ ----(Δ64 bp)----ATCGGGCGTTCGAC |
| #44 | GGAATG CT----(Δ279 bp)----ACAGACCAT |
| #46 | GGAATG CTG$_{5}$ C----(Δ260 bp)----ATCACAGACCAT |
| #17 | CCTTGTAGCCGGGA----(Δ281 bp)----TCACAGACCATTTC |
| #19 | TCCTTGTAGCCGGG----(Δ312 bp)----CTGAGGCCGACCTG |
| #31 | TCGTCCTTGTAGCC----(Δ295 bp)----ATTTCTTTCTTTCG |
| #4 #7 | (δ20)TTTGTCACTCATTATCCTATTACATTATCAATCCTT----(Δ3050 bp)----TCATTATCCTATTACATTATCAATCCTT GCATTTCAGCTTCC(δ18) |
| #39 | (δ20)TAATATTAGGTATACAGAATATACTAG----(Δ2837 bp)----AGGTATACAGAATATACTAG AAGTTCTCCTCGAG(δ18) |

***rad52*Δ junctions**

| | |
|---|---|
| Parental allele | GGAATG CTG$_{80}$ CTGGGGGGATCACAGACCAT |
| #B10 #B12 | GGAATG CTG CTGGGGGGATCACAGACCAT |
| #A3 | GGAATG CTG$_{3}$ CTGGGGGGATCACAGACCAT |
| #C8 | GGAATG CTG$_{4}$ CTGGGGGGATCACAGACCAT |
| #C5 | GGAATG CTG$_{5}$ CTGGGGGGATCACAGACCAT |
| #H4 | GGAATG CTG$_{8}$ CTG----(Δ262 bp)----CTGCAGCGAA |
| #A1 | GGAATG C---------(Δ255 bp)-------------CCAT |
| #A2 | GTCGTCCTTG----(Δ291 bp)---TGAGGCCGAC |
| #A11 | AAGCTGGTCA----(Δ579 bp)---CGACCTGCAG |
| #B6 | GGAATG----------(Δ252 bp)-------CAGACCAT |
| #B7 | GGAATG CTG$_{3}$ CT-(Δ276 bp)-ACCTGCAGCG |
| #C9 | GTCGTCCTTG----(Δ305 bp)---AGCGAAATCT |
| #C4 | (δ20)ATCCCACCAATTATC-(Δ3021 bp)- CAATTATCAAAAAAT(δ18) |
| #C7 | (δ20)TTATCTCAAAATTCAC-(Δ3029 bp)-AAAATTCACCTTTCTT (δ18) |

***sae2*Δ junctions**

| | |
|---|---|
| Normal allele | GGAATG CTG$_{77}$ CTGGGGGGATCACA |
| #A8 #D2 #D12 | GGAATG CTG$_{3}$ CTGGGGGGATCACA |
| #D3 | GGAATG CTG$_{72}$ (C)TGCTGGGGGGATCACA |
| #A2 | GGAATG CTG$_{77}$ CGCTGGGGGGATCACA |
| #A4 | GGAATG CTG$_{77}$ C(T)GGGGGGGGATCACA |
| #A5 | GGAATG CTG$_{77}$ CCTGCTGGGGGGATCACA |
| #A6 | GGAATG CTG$_{77}$ CTGCGCCTGGGGGGATCACA |
| #A11 | GGAATG CTG$_{77}$ CTTGGTGGGGGGATCACA |
| #A12 | GGAATG CTG$_{77}$ CGC(T)GGGGGGGATCACA |
| #B5 | GGAATG CTG$_{77}$ CTGCTGCGCTGGGGGGATCACA |
| #E2 | GGAATG CTG$_{18}$ CCTGCTGGGGGGATCACA |
| #B2 | (δ18)ATAATGTAATAGGATAATGA-----------(Δ3027 bp)-----------TAATAGGATAATGAGTGACA(δ20) |

***dnl4Δ* junctions**

| | |
|---|---|
| **Parental allele** | **GGAATG CTG$_{80}$ CTGGGGGGATCACA** |
| **#A9** | **(δ20)AAGACTTAATATTTAGGTATACAGAATATACTAGGAG----(Δ3029 bp)----CAGGTATACAGAATATACTAGAGTTCTCCTCGA(δ18)** |
| **#B2** | **(δ20)TTTCATTAGACTTTGTTACTGTT----(Δ3050 bp)----ACAGTTCCTCGTATCTTATGTCATCG(δ18)** |
| **#B7 #B9 #D3** | **(δ20)ATGTTTGGAAGAAAGGTGAATTTT----(Δ3052 bp)----GGTGAATTTTGAGATAATTGGTGGG(δ18)** |
| **#B12** | **(δ20)GTAAAGACTTAATATTAGGTATACAGAATATACTAG----(Δ3053 bp)----AGGTATACAGAATATACTAGAAGTTCTCCTCGAGGA(δ18)** |
| **#C10 #C12** | **(δ20)CTTATGTTATCTTCTTACACAGTAT----(Δ3049 bp)----ACACCGTATATGATAATATATTGGT(δ18)** |
| **#E8 #E9 etc.** | **(δ20)TGATACTTTGTCACTCATTATCCTATTACATTATCAA----(Δ3051 bp)----TCATTATCCTATTACATTATCAATCCTTGCATTTCAG(δ18)** |
| **#C9 #E11** | **(δ20)TGTTGGTAAAGACTTAATATTAGGTATACAGAATATACTAG----(Δ3054 bp)----TAATATCAGGTATACAGAATATACTAGAAGTTCTCCTCGAG (δ20)** |
| **#B5** | **(δ20)AAAGGTGAATTTTTTGATAATTG----(Δ2798 bp) +153 bp Tyδ19 chr.10----ATCCTATTACATTATCAATCCTT(δ19)** |
| **#D11** | **(δ20)TTGCTGGGATTCCATTGTTG----(Δ3147 bp) +94 bp Ty1δ21 chr.7----ATTACGATTATTCCTCATTC(δ18)** |

***sae2Δ dnl4Δ* junctions**

| | |
|---|---|
| **Parental allele** | **GGAATG CTG$_{81}$ CTGGGGGGATCACA** |
| **#B11** | **GGAATG CTG$_{81}$ CTGGGGGGATCACA** |
| **#B8 #C12 etc.** | **GGAATG CTG$_{80}$ CTGGGGGGATCACA** |
| **#C9 #D1 etc.** | **GGAATG CTG$_{79}$ CTGGGGGGATCACA** |
| **#A2** | **GGAATG CTG$_{3}$ CTGGGGGGATCACA** |
| **#A3** | **(δ20)GTTTCTCAATCCTTATGT----(Δ3050 bp)----CTTATGTCATCGTCTAACACCGTATATGA(δ18)** |
| **#A8** | **(δ20)GTTTCTCAATCCTTATGTTATC----(Δ3050 bp)----CTTATGTCATCGTCTAACACCGTAT(δ18)** |
| **#C4 #C5 etc.** | **(δ20)ATTACATTATCAATCCTTGCATTT----(Δ3051 bp)----CAATCCTTGCATTTCAGCTTCCTC(δ18)** |
| **#B6** | **(δ20)GTTTGGAAGAAAGGTGAATTTT----(Δ3052 bp)----AAAGGTGAATTTTGAGATAATT(δ18)** |
| **#B7 #C3** | **(δ20)ATATTAGGTATACAGAATATACTAG----(Δ3053 bp)----AGGTATACAGAATATACTAGAAGTT(δ18)** |
| **#A4** | **(δ20)GTATACAGAATATACTAG----(Δ3226 bp) +176 bp Tyδ12 chr.5----TCTAACACCGTATATGAT(δ19)** |
| **#C8** | **(δ20)TTCTTTGATA--(Δ25 bp) +23 bp Tyδ16 chr.10--TTATCAATCCTTGCATTT----(Δ3033 bp)----TTATCAATCCTTGCATTTCAGCTTCCTC(δ18)** |

**Wild-type e*Sp*Cas9 junctions (guideRNA #1)**

| | |
|---|---|
| Parental allele | GGAATG CTG$_{78}$ CTGGGGGGATCACAGACCATTTCTTT |
| C4# | TCGTCCTTGTAGCCGGG------(Δ280 bp)-----GGGGATCACAGACCATT |
| D12# | GGAATG CTG$_{6}$C--------(Δ2541 bp)--------GATGGAAATCATTATTGTAGT(δ18) |
| B4# B5# | (δ20)TATCTTCTTACACAGTAT------(Δ3050 bp)------GTATATGATAATATATTG(δ18) |
| A3# A5# etc. | (δ20)ACTTTGTCACTCATTATCCTATTACATTATCAA------(Δ3051 bp)------TCATTATCCTATTACATTATCAATCCTTGCATTT(δ18) |
| B1# | (δ20)TTCCATTGTTGGTAAAGACT(A)TAATAT------(Δ3053 bp)------ATTGTTGGTAAAGGCTATAATATCAGGTATACAGAAT(δ18) |

**Wild-type e*Sp*Cas9 junctions (guideRNA #2)**

| | |
|---|---|
| Parental allele | GGAATG CTG$_{78}$ CTGGGGGGATCACAGACCATTTCTTT |
| #A8 | GGAATG CTG$_{17}$ TTG CTG$_{5}$ CTGGGGGGATCACAGACCA |
| #A12 | GGAATG CTG$_{66}$ ----------G CTGGGGGGATCACAGACCA |
| #C4 | GGAATG CTG$_{25}$ (C)TTTGGGGGGATCACAGACCATTTCTTT |
| #A3 | GGAATG CTG$_{13}$ CTG--------(Δ238 bp)--------GATCACA |
| #A4 | TAGCACCACGCC-----(Δ1342 bp)-----CCCCCGGGAGAT |
| #A6 | TAATTTGTCAGT-----(Δ604 bp)------CTTGCAAGTTGA |
| #B6 | TCTCGGTAGCCA------(Δ401 bp)------CGTTCGACTCGC |
| #B9 | (δ20)ATAATATAATAGTAACATGAA------(Δ2404 bp)------TGTTGGAATAAAAATCAACTATCATC(δ19) |
| #A2 #A5 etc. | (δ20)TATCAATCCTTGCATTT-------(Δ3017 bp)-------TATCAATCCTTGCATTT(δ18) |
| #C9 | (δ20)ATTGTTGGTAAAG-------(Δ3039 bp)-------ATTGTTGGTAAAG(δ18) |
| #E11 | (δ20)ACTCATTATCCTATTACATTATCAATCCTTGCATTT------(Δ3051 bp)------TCCTTGCATTTCAGCTTCCTCTAAC(δ18) |
| #D10 #E3 #F7 | (δ20)TTAATATTAGGTATACAGAATATACTAG------(Δ3053 bp)------AGGTATACAGAATATACTAGAAGTTCTCCTCGAG(δ18) |
| #F8 | (δ20) TTCCTTCTTTGATA-------(Δ3112 bp)+59 bp Tyδ16 chr.10-------ATGACAGTTCCTCG(δ18) |

**Wild-type *Sp*Cas9 junctions (at the *ARG2* locus)**

| | |
|---|---|
| Parental allele | GGAATG CTG$_{60}$ CTGGGGGGATCACAGACCATTTCTTT |
| #B10 | GGAATG CTG$_{72}$ CTGGGGGGATCACAGACCATTTCTTT |
| #A2 #A3 etc. | AGGCGCAAGACTTCAA----(Δ388 bp +TAGGGATAACAGGGTAAT)----CGAAATCTTGAGATCG |
| #B5 | CATCTAGAGTCGTCCTTGTAGCC--(Δ325 bp)--TTTCTTTCTTTCGGCCAGGCTG |
| #A8 #B11 | TCTCCACAGCAGCAGC-------(*YAK1-ARG2*)-------ATCACAGACCATTTCT |

**Supplemental Figure 1**

Mosbach *et al.*
