## Supplemental Figure 2 for "Resection and repair of a Cas9 double-strand break at CTG trinucleotide repeats induces local and extensive chromosomal rearrangements"

**A**

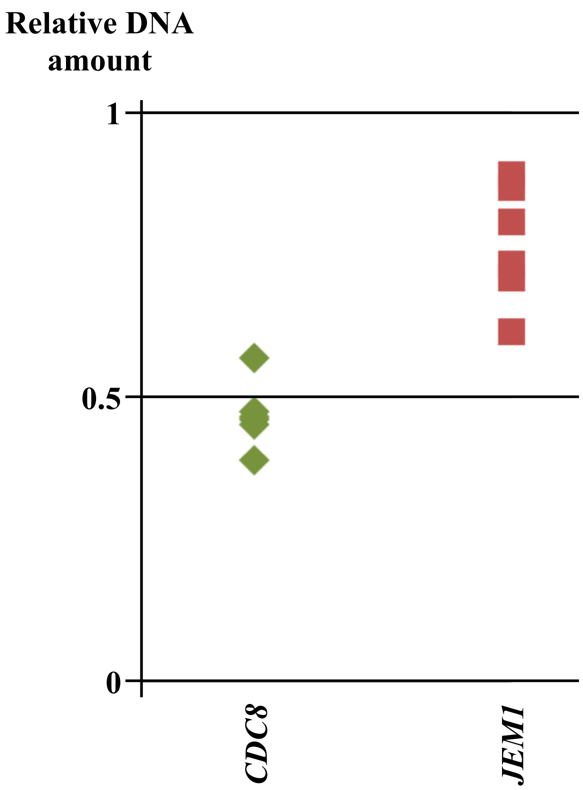


**B**

| Diploids | | | | | | | | Haploids | | | |
|---|---|---|---|---|---|---|---|---|---|---|---|
| Cas9 repressed | | | | Cas9 expressed | | | | Cas9 expressed | | | |
| Clone | Substit. | Indel | Indel micro. | Clone | Substit. | Indel | Indel micro. | Clone | Substit. | Indel | Indel micro. |
| 1 | - | - | - | 1 | - | - | 1 | 1 | - | - | - |
| 2 | 1 | - | - | 2 | - | - | 1 | 2 | 1 | - | - |
| 3 | 1 | - | - | 3 | - | - | - | 3 | - | - | - |
| 4 | - | - | - | 4 | - | - | - | 4 | - | - | - |
| 5 | - | - | - | 5 | - | - | - | 5 | - | - | - |
| **Total** | **2** | **0** | **0** | 6 | - | - | - | 6 | 1 | - | - |
| | | | | 7 | - | - | - | 7 | 1 | - | - |
| | | | | 8 | - | - | - | 8 | 2 | - | - |
| | | | | 9 | - | - | - | 9 | 2 | - | - |
| | | | | 10 | - | - | - | 10 | 1 | - | - |
| | | | | 11 | - | - | - | **Total** | **8** | **0** | **0** |
| | | | | 12 | - | - | - | | | | |
| | | | | 13 | - | - | - | | | | |
| | | | | 14 | - | - | - | | | | |
| | | | | 15 | 1 | - | - | | | | |
| | | | | 16 | - | 1* | - | | | | |
| | | | | 17 | - | - | - | | | | |
| | | | | 18 | 2 | - | - | | | | |
| | | | | **Total** | **3** | **1** | **2** | | | | |

**Supplemental Figure 2**
**Mosbach *et al.***
