## Supplemental Table 1 for "Resection and repair of a Cas9 double-strand break at CTG trinucleotide repeats induces local and extensive chromosomal rearrangements"

**Supplemental Table 1: list of strains used in the present study**

| **Strain** | **Genotype** | **Origin** |
| --- | --- | --- |
| GFY6162-3D | *MAT*α *ura3*Δ851 *leu2*Δ1 *his3*Δ200 *trp1*Δ63 *ade2*-opal *SUP4*::(CAG)255 | (Richard et al., 2003) |
| VMY350 | *MAT*α *ura3*Δ851 *leu2*Δ1 *his3*Δ200 *trp1*Δ63 *ade2*-opal *SUP4*::(CAG)80 *rad50*Δ::KANMX4 | GFY6162-3D |
| VMY650 | *MAT*a *ura3*Δ851 *leu2*Δ1 *his3*Δ200 *trp1*Δ63 *ade2*-opal *SUP4*::(CAG)80 *rad50*Δ::KANMX4 | VMY350^(1)^ |
| VMY352 | *MAT*α *ura3*Δ851 *leu2*Δ1 *his3*Δ200 *trp1*Δ63 *ade2*-opal *SUP4*::(CAG)80 *rad52*Δ::KANMX4 | GFY6162-3D |
| VMY452 | *MAT*a *ura3*Δ851 *leu2*Δ1 *his3*Δ200 *trp1*Δ63 *ade2*-opal *SUP4*::(CAG)80 *rad52*Δ::KANMX4 | VMY352^(2)^ |
| VMY104 | *MAT*α *ura3*Δ851 *leu2*Δ1 *his3*Δ200 *trp1*Δ63 *ade2*-opal *SUP4*::(CAG)80 *dnl4*Δ::KANMX4 | GFY6162-3D |
| GFY199 | *MAT*α *ura3*Δ851 *leu2*Δ1 *his3*Δ200 *trp1*Δ63 *ade2*-opal *SUP4*::(CAG)80 *sae2*Δ::KANMX4 | GFY6162-3D |
| GFY205 | *MAT*α *ura3*Δ851 *leu2*Δ1 *his3*Δ200 *trp1*Δ63 *ade2*-opal *SUP4*::(CAG)80 *sae2*Δ::*HIS3* *dnl4*Δ::KANMX4 | VMY104 |
| GFY117 | *MAT*a *ura3*Δ851 *leu2*Δ1 *his3*Δ200 *trp1*Δ63 *ade2*-opal *SUP4*-opa*l*::I-*Sce*Ics *arg2*Δ216::*SUP4*::(CAG)_98_-*TRP1* | (Richard et al., 2003) |

^(1)^ VMY650 was built by switching VMY350 mating-type with the pJH132 HO-expressing plasmid (Holmes and Haber, 1999).

^(2)^ VMY452 was built by switching VMY352 mating-type with the pJH132 plasmid.
