## Supplemental Table 2 for "Resection and repair of a Cas9 double-strand break at CTG trinucleotide repeats induces local and extensive chromosomal rearrangements"

**Supplemental Table 2: list and sequence of primers used in this study**

| **Name** | **Oligonucleotide sequence** |
| --- | --- |
| VMS1  VMAS1 | 5’-GATGCTGCCCATGCTATAGA-3’  5’-CCCACTGACAACAACATTGG-3’ |
| VMS2  VMAS2 | 5’-TTCCGAGACGCAAATCACAG-3’  5’-GTCCTCAACGTCGTTCTCAA-3’ |
| VMS3  VMAS3 | 5’-TCTGTGCCGCAATATACCTC-3’  5’-TTGGACGACCCTGTGTTGAA-3’ |
| VMS4  VMAS4 | 5’-AATTGTCGAACAACACGGCG-3’  5’-TGAGAGACGTCGTTGTAACC-3’ |
| VMS6  VMAS6 | 5’-TGGAGGAAGGACGAAACACT-3’  5’-TGAGAGTCATCCATGTGCCA-3’ |
| SAE2up  SAE2down | 5’-CCAGTAATTGACGATGCGGAAG-3’  5’-GCAGAACTATGTGAGTGATGACC-3’ |
| Su47  Su48 | 5’-AATTGAATATATTCTGCTCTTGAATTAGATatgtg-3’  5’-ATAGCTATGGAAAATAACGCAGCAGC-3’ |
| Su23  Su42 | 5’-GGGGTACCTTATGACTTCTTTTTCGCCTACTA-3’  5’- GTTTAGAAGCGTACTTCTTGGG-3’ |
| VMS20  VMAS20 | 5’-AGATTTATCCTGCGAGTTTG-3’  5’-TGACTGCTGACGAGTTAAG-3’ |
| VMS21  VMAS21 | 5’-AGCACTACATACGTGACTTT-3’  5’-GACTTCTTTGGAGTACTGTGA-3’ |
| VMS22  VMAS22 | 5’-TGTGCATTGGTAACAAAGGG-3’  5’-AGAAGCCGTTATTCTCTTCTT-3’ |
| VMS23  VMAS23 | 5’-TCCAATAAACATCTCGTCCA-3’  5’-CTAGATGCATTTTTCGGCAA-3’ |
| JEM1f  JEM1r | 5’-TGTGATTTGGCTGAGTTACAACG-3’  5’-AACTGCCCAGCGATCCATT-3’ |
| Chromo4_f  Chromo4_r | 5’-TTCCACGGGTAGGTACGACT-3’  5’-TTTCAAGGACCCTCGTTCGG-3’ |
| LP400 | 5'- CCCCGGATTCTAGAACTAGTGGATCCCCCGGGAAAAAAATGGACTATAAGGACCACGACG-3' |
| LP401 | 5'-TAAGCGTGACATAACTAATTACATGACTCGAGAAGAGATTACTTTTTCTTTTTTGCCTGGCC-3' |
| SNR52Left | 5'-gtctaaggcgcctgattcaagaaatatcttgaccgcagttAACtctttgaaaagataatgta-3' |
| SNR52Right | 5'- gaactcttgcatcttacgatacctgagtattcccacagttAACAGACATAAAAAACAAAAAA-3' |
| gRNA#1 | 5'-GCTGCTGCTGCTGCTGCTGC-3' |
| gRNA#2 | 5'-TGCTGCTGCTGCTGCTGCTG-3' |
